# Near-Atomic Cryo-EM Imaging of a Small Protein Displayed on a Designed Scaffolding System

**DOI:** 10.1101/212233

**Authors:** Yuxi Liu, Shane Gonen, Tamir Gonen, Todd O. Yeates

## Abstract

Current single particle electron cryo-microscopy (cryo-EM) techniques can produce images of large protein assemblies and macromolecular complexes at atomic level detail without the need for crystal growth. However, proteins of smaller size, typical of those found throughout the cell, are not presently amenable to detailed structural elucidation by cryo-EM. Here we use protein design to create a modular, symmetrical scaffolding system to make protein molecules of typical size amenable to cryo-EM. Using a rigid continuous alpha-helical linker, we connect a small 17 kDa protein (DARPin) to a protein subunit that was designed to self-assemble into a cage with cubic symmetry. We show that the resulting construct is amenable to structural analysis by single particle cryo-EM, allowing us to identify and solve the structure of the attached small protein at near-atomic detail, ranging from 3.5 to 5 Å resolution. The result demonstrates that proteins considerably smaller than the theoretical limit of 50 kDa for cryo-EM can be visualized clearly when arrayed in a rigid fashion on a symmetric designed protein scaffold. Furthermore, because the amino acid sequence of a DARPin can be chosen to confer tight binding to various other protein or nucleic acid molecules, the system provides a future route for imaging diverse macromolecules, potentially broadening the application of cryoEM to proteins of typical size in the cell.

**Significance statement:** *New electron microscopy methods are making it possible to view the structures of large proteins and nucleic acid complexes at atomic detail, but the methods are difficult to apply to molecules smaller than about 50 kDa, which is larger than the size of the average protein in the cell. The present work demonstrates that a protein much smaller than that limit can be successfully visualized when it is attached to a large protein scaffold designed to hold 12 copies of the attached protein in symmetric and rigidly defined orientations. The small protein chosen for attachment and visualization can be modified to bind to other diverse proteins, opening up a new avenue for imaging cellular proteins by cryo-EM.*

## INTRODUCTION

Recent advancements have brought single particle electron microscopy techniques to the forefront of structural biology (1–3). In favorable cases, three-dimensional cryo-EM image reconstruction methods are able to produce structures of macromolecular complexes at atomic level detail (4–9). In such studies, very large macromolecular assemblies offer important advantages in signal processing and imaging, and this advantage is enhanced in systems that are highly symmetric – e.g. composed of many repeating copies of one or a few protein building blocks. For those reasons, viral capsids are quintessential examples for favorable cryo-EM reconstruction. At the other end of the spectrum, however, individual protein molecules of typical size (e.g. 50 kDa or smaller), which lack the aforementioned advantages, remain extremely difficult to visualize at atomic detail by electron microscopy. This critical size limitation represents a singular impediment to the universal application of electron microscopy for elucidating structures of most proteins in the human genome.

Recent studies have shown that small proteins can be computationally re-designed so that multiple copies of the protein subunit will self-assemble into large, symmetric cages with shapes resembling regular geometric solids: e.g., a tetrahedron, cube, or icosahedron (10–16). The structures resulting from some of these designed assembly approaches have sufficiently large mass and high symmetry that they can be analyzed readily by cryo-EM. However, current methods for designing protein assemblies are laborious and unpredictable, often requiring substantial trial-and-error experiments and prior structural knowledge to achieve success. Those challenges have made it impractical to take a given target protein of interest, whose structure may not be known, and engineer it to assemble into a large symmetric assembly that would be amenable to cryo-EM.

It would advance cryo-EM applications tremendously if it were possible to easily attach a protein of interest to a symmetric scaffold in a rigid way, so that many copies of the target protein would be displayed in well-defined, symmetric orientations. Being able to turn a given protein into its own kind of capsid structure would confer on it the features of large size and symmetry that are critically advantageous for cryo-EM imaging. In the present study we demonstrate a route towards that ultimate goal (Fig. 1A).

**Figure 1.**
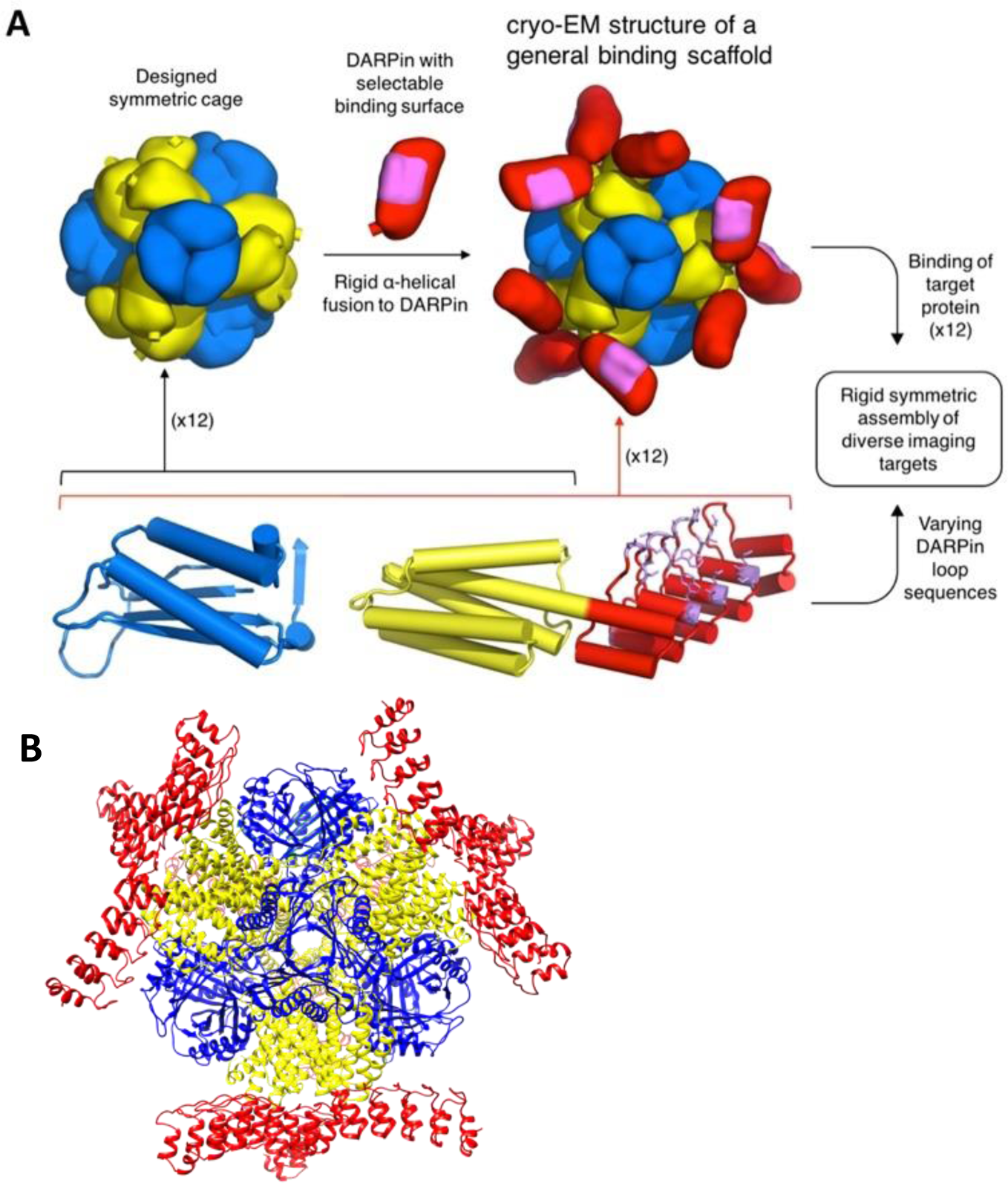
A molecular scaffolding system for modular display of macromolecules for cryo-EM imaging. A. Schematic diagram for a scaffolding system built upon a designed symmetric protein cage; the example shown is a tetrahedrally symmetric cage with 24 subunits in a_12_b_12_ stoichiometry (A subunits in yellow and B subunits in blue). At least one of the subunits needs to have an α-helical terminus (cylinder). The α-helical termini of the cage subunit (yellow) and the DARPin subunit (red) can be joined in a rigid fashion through genetic fusion, forming a general binding scaffold. The cryo-EM structure of a binding scaffold is solved in this study. The DARPin subunit contains variable loops (highlighted in pink) whose amino acid sequence can be selected to confer binding to a wide range of specific macromolecules of interest. In principle, binding a macromolecule of interest to the designed scaffold results in the symmetric display of 12 copies of the molecule. B. Detailed view of the specific scaffold, DARP14, that was designed and characterized in this study, with subunits colored as in panel A.

## RESULTS

For designing a modular cryo-EM scaffold, we took as a starting point a set of protein cages designed by King et al.(14), specifically those built from 24 subunits, four trimers of two different subunit types. These assemble with the different trimer types sitting at alternating corners of a cube, in arrangements that obey tetrahedral symmetry. In the current study, we focused our attention on designed protein cages where one or both component subunit types contain at least one alpha helical terminus. Through further design, we extended the alpha helical terminus of the cage protein by genetic fusion to join the alpha helical terminus of a small protein target of only ~17 kDa known as DARPin (Designed Ankyrin Repeat Protein) (17). This design element, fusing two proteins with terminal helices, is intended to create a semi-rigid and geometrically predictable helical connection spanning between two proteins that would otherwise be flexibly joined (Fig. 1A). This idea was developed by Padilla et al. and subsequently expanded upon and validated in various contexts (10, 11, 18, 19).

The choice of a DARPin as the first fusion partner to the cage is critical, as DARPins have been developed as a general platform for binding other protein molecules. Through genetic selection techniques, amino acid sequence changes in loop regions of the DARPin protein can be identified for conferring tight binding to various target proteins of interest (20–23). In addition, their largely alpha-helical nature makes DARPins suitable for fusion to other proteins by the continuous alpha helical fusion approach. Taken together, the essence of our scaffolding system is that a rigid protein cage forming a core structure will present (as genetic fusions) 12 rigid and symmetrically disposed DARPin proteins projecting outward (Fig. 1). In the future, loop sequences specific for binding some other target protein can be readily exchanged into the basic DARPin structure, thereby enabling the facile capsid-like assembly of varied target proteins. Importantly, this strategy ultimately circumvents the need to perform engineering experiments on future targets themselves by restricting design efforts to the protein cage and its fused DARPin.

We experimentally tested several variations in the amino acid sequence and length of the helical connection between the cage subunit and the DARPin based on computationally generated fusion models (see Methods and Supporting Information Text). We devoted our efforts to specific design choices that disposed the DARPin binding surfaces in highly accessible orientations for subsequent utility in binding cognate target proteins. Among the designs investigated, five could be purified in soluble form from a bacterial overexpression system and were shown to self-assemble into structures of the expected size and shape by negative-stain electron microscopy (Supporting Information Fig. 1).

Next, we pursued a full structural elucidation for one of the scaffold designs, referred to here as DARP14, by 3D cryo-EM reconstruction (Fig. 2 & 3, and Supporting Information Table 1). DARP14 was imaged on a Titan Krios using a K2 direct electron detector (see Methods). A total of 3665 movies were recorded for motion correction and after reference-free 2D classification 229,953 particles were selected for 3D analysis. In the raw cryo-EM images, the core of the protein cage was discernible but the individual DARPin components appeared weaker or were practically invisible (Fig. 2A). This was expected, and further reinforced the well-known challenge of imaging small protein molecules on their own. Subsequent 2-D class averages and 3-D reconstruction showed the powerful advantage of being able to locate and apply symmetry-averaging to the smaller DARPin components when displayed on the engineered scaffold (Fig. 2 & 3). A 3-D reconstruction of the cage based on a subset of 34,650 particles produced an image with an overall resolution of ~3 Å (Fig. 2C and Supporting Information Table 1). The side chains of the amino acids in the core of the cage are clearly discernable in the resulting density maps and are consistent with the designed protein. This is the first example of an atomic resolution structure determined by single particle cryo-EM of a designed protein cage.

**Figure 2.**
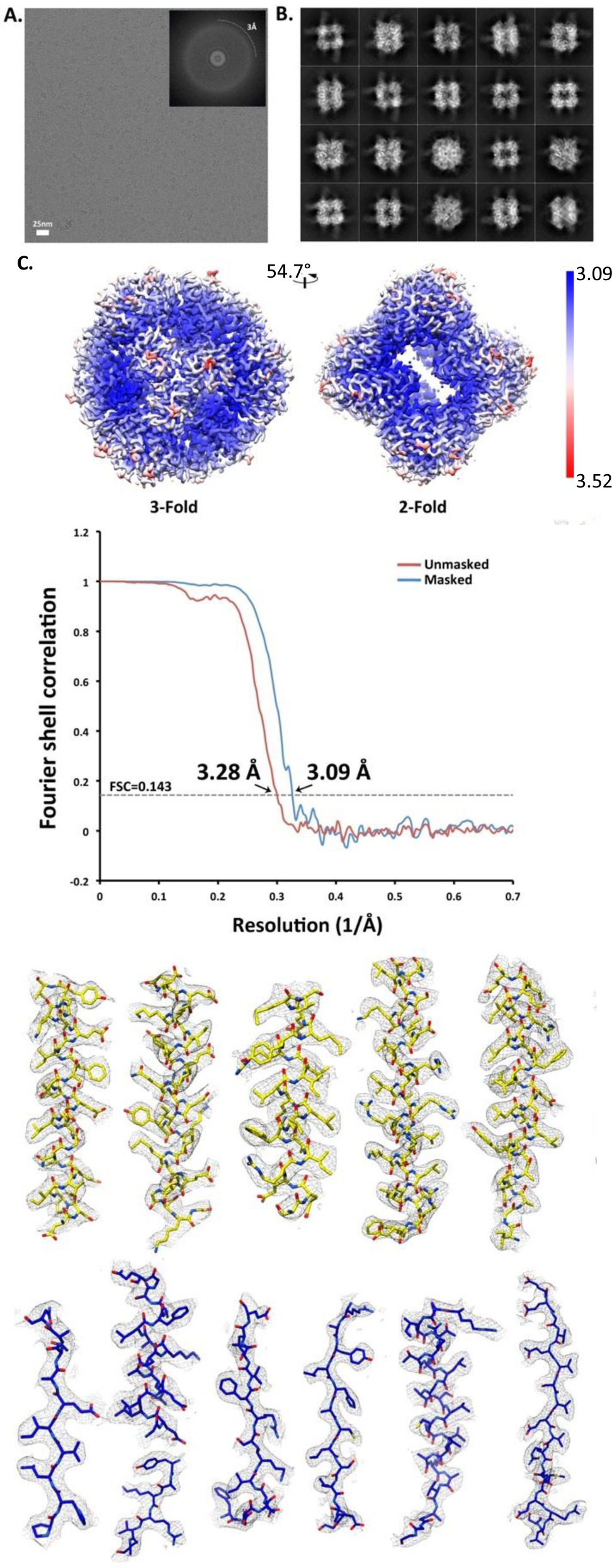
Cryo-EM structure of DARP14 symmetric cage core. A. Representative motion-corrected cryo-electron micrograph of DARP14. (inset) Fourier transformation showing visible thon rings to ~3Å. B. Reference-free 2D class averages highlighting good alignment of the cage and clear density for fused 17kDa DARPins. C. Overview of a ~3.1 Å reconstruction of the cage core. Top: Representations of unfiltered local resolution viewing down the 3-fold and 2-fold symmetry axes. Middle: Fourier shell correlation (FSC) curves of unmasked and masked reconstructions. Bottom: refined model fit into density for subunit A (yellow) and subunit B (blue) of the cage core. All secondary structure elements are represented along with selected loop regions.

Importantly, much of the attached DARPin was also visible in the 2D class images (Fig 3A) and reconstructions. To account for the possibility of slight variations in the orientations of the attached DARPins, which would compromise their resolution, we applied subsequent classification to 183,753 particles after masking out the B type subunit of the cage in order to focus on the DARPin component (see Methods). This substantially improved the structural details visible for the DARPin, resulting in a ~3.5 Å resolution structure overall (Fig 3B-C) and allowing us to clearly model the helical secondary structural elements within the density (Fig. 3D) in configurations consistent with the known crystal structure of the DARPin (PDB 3ZU7) (Supporting Information Table 1).

**Figure 3.**
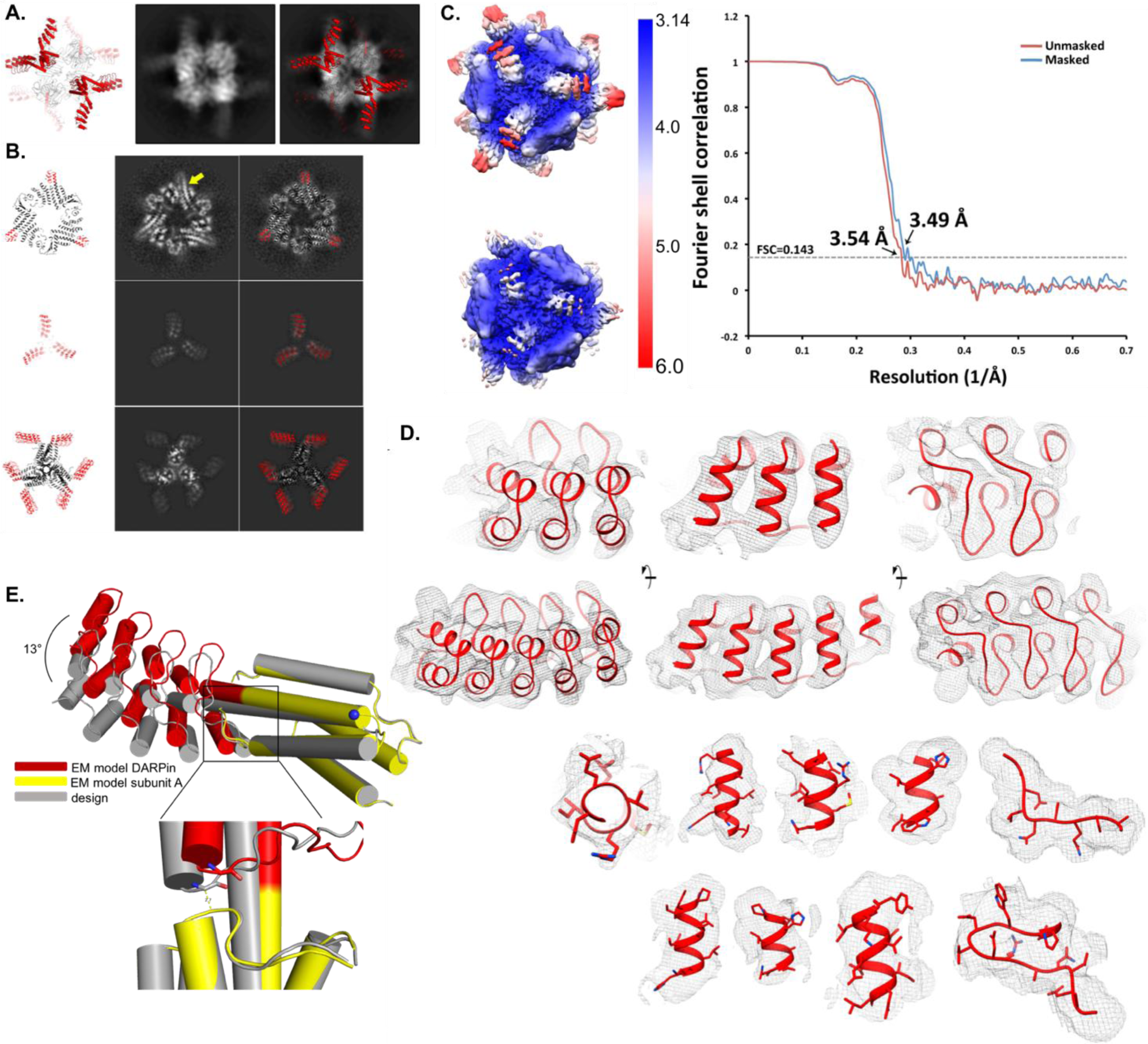
Cryo-EM reconstruction of DARPin displayed on the symmetric cage. A. Comparison of the DARP14 design (DARPins in red and cage subunits A and B in black and white) to one 2D class average with overlay (right) highlighting density for DARPin helices protruding from the cage. B. Three comparisons of the calculated model and slices of reconstructions. Top: Focus on the extended helix where DARPins are fused (yellow arrow). Middle: Top view showing the DARPin arms and clear density for each helical repeat. Bottom: Side-slice. C. Local resolution of unfiltered ~3.5Å reconstruction where the subunit A and fused DARPin were masked during refinement for higher resolution of those areas. Left top: Low contouring level to show entire reconstruction. Left bottom: Higher contouring level highlighting the near-atomic detail of DARPin repeats. Right: FSC of unmasked and masked DARPin reconstructions. D. Highlight of DARPin density in different regions with the fitted model. Top two rows: High sigma level highlighting DARPin helical repeats 1-3 (top) and lower sigma level highlighting all 5 helical DARPin repeats as a top view (left), side view (middle) and bottom view (right). Views related by 90° rotations. Bottom two rows: Density fit of DARPin model from various helices (including one top view of helix 2) and two views of loop regions where the amino acid sequence for the DARPin would be varied for binding to cognate target molecules. E. Comparison between the computational design for DARP14 and the cryo-EM density-fitted model, showing a small displacement of the fitted model from the design. The designed DARP14 and the cryo-EM model were aligned on their A subunits (which is named chain B in PDB 4NWP). Top: there is a ~13° rotation around an axis going through the view of plane at the blue dot between the design and the EM model. Bottom: zoom in at Gly 187 in the first turn on the DARPin, which is in close proximity to Gly108 and Thr 109 from subunit A in the design. That interaction appears to stabilize the DARPin in its observed orientation.

The DARPin protein we attached to the cage is comprised of five repeats of a common structural motif (the ankyrin repeat). In our final 3-D reconstruction of the DARPin, the first four repeats could be resolved at near-atomic detail, with the local resolution worsening from 3.5 Å to 5 Å toward the tip of the structure (Fig. 3C). This worsening of the resolution toward the tips of cryoEM structures has been observed in other cryoEM studies (24). Moreover, we suspect that the fifth DARPin repeat in our designed scaffold may be flexible and partially unwound, further contributing to its weakness in the final image. Consistent with this explanation, the thermal vibration parameters (B factors) in previous crystal structures of DARPins are higher for this region of the protein (25, 26) (Supporting Information Fig. 2). We note that this tendency toward terminal unwinding is not an impediment to forming a well-ordered complex between a DARPin and its cognate target, as demonstrated in multiple previous crystal structures where the DARPin and its target are well-ordered when bound together (21, 27–29). Notwithstanding the loss of resolution at the end of the attached DARPin, the final result represents the first example of a small protein being visualized at near-atomic resolution by a cryo-EM scaffolding approach.

## DISCUSSION

Our analysis demonstrates that the alpha helical fusion scheme used here provides a connection between the symmetric cage and the DARPin that is rigid enough to enable near atomic-resolution imaging. This is a critical result as it was not known in advance whether the alpha helical fusion would hold the DARPin in a sufficiently ordered configuration. The ordered nature of the DARPin was evident in preliminary 2D averaging (Fig. 3A) even before 3D reconstruction and application of symmetry to optimize the imaging of the cage. In comparing our final structure to the initial computational model, a minor reorientation of the DAPRin component (by approximately 13 degrees) is evident (Fig. 3E). Among the several designs we explored (Supporting Information Fig. 1), the structure of the one analyzed here appears to be influenced, beyond our designed continuous alpha-helical fusion, by a few additional atomic contacts between the DARPin and the cage subunits. These contacts likely help stabilize the DARPin in a well-defined orientation on the scaffolding cage. The relatively high orientational rigidity we obtained for the DARPin promises good prospects for similarly rigid attachment of other proteins to the DARPin for their visualization in subsequent studies. Past studies of DARPin complexes indicate stable and rigid binding to their cognate protein targets (*21*, *26*–*28*).

Our results emphasize two major points. First, the DARPin component is a small protein (17 kDa) whose separate structure would otherwise be impossible to resolve by single particle cryo-EM methods. Yet it can be visualized in near atomic detail when its image is reconstructed in the context of rigid assembly on a large symmetric protein cage. Recent work by Coscia et al. (30) was able to image a larger (40 kDa) target protein fused to a natural protein scaffold at lower resolution (local resolution between 6 and 10 Å) and only after extensive biochemical analysis and optimization of linker lengths. We show here that a rational design of a continuous alpha helical attachment to a cubically symmetric designed protein cage can provide the rigidity required to achieve near atomic resolutions even for a small 17 kDa attached target protein. Since our present scaffold was the best among only a relatively small number of candidates investigated, it is likely that further design efforts could improve the degree of rigidity, making it possible to reach an even better spatial resolution. Second, our development of a DARPin as the fused protein component introduces a critical element of modularity. Building on this system, the challenging molecular engineering required to create symmetric architectures will not need to be repeated for each application to a new target protein to be imaged; in principle, no modification to a future target protein is required. In the future, novel target proteins could be attached to the DARPin by identifying DARPin loop sequences that bind the target protein, as has been demonstrated in other studies (22, 23). Notably, each distinct scaffolding system that might be created by variations on the design theme developed here, and each different DARPin sequence selected for binding to a given target protein, will provide a distinct opportunity for obtaining a high-resolution structure of a target protein. Further developments on this scaffolding approach should ultimately enable the facile imaging of large numbers of cellular proteins whose structures have previously been beyond the reach of cryoEM.

## MATERIALS AND METHODS

### Computational α-helix fusion methods

Computational α-helix fusion models were generated similarly to our previous work (10, 11). As a test case for fusing to a protein cage, we used a DARPin whose sequence was selected to bind to the extracellular signal-regulated kinase 2 (ERK2), and whose structure in complex with its cognate partner is known (PDB 3ZU7). In choosing a protein cage as the fusion partner, we restricted our attention to those that have a protein with a terminal α-helix at least six amino acids long and with no more than ten unstructured amino acids beyond it. The set of protein cages that satisfied this criteria included six protein assemblies designed in previous work (11, 14, 15, 31, 32). Next, we tested the feasibility of pair-wise joining between the protein cage subunit and the DARPin subunit. To do so, we first aligned an ideal α-helix to the last six helical residues on the cage subunit. Then we aligned the DARPin terminal helix to the ideal α-helix. The aligned position of the DARPin on the ideal α-helix was slid one residue at a time. The range of sliding was from a six-residue overlap to a 15-residue insertion relative to the helical termini of the DARPin and the cage subunit. We inspected the models at each aligned position and removed those with excessive clashes. If the fusion model had overlapping helical termini, the amino acid sequence within the overlap was chosen to maintain good native contacts within each subunit. If the fusion model required an insertion between the helical termini, ER/K-rich helix segments (19, 33) were used. The final set of experimentally tested models were chosen to give different DARPin orientations relative to the cage subunit while providing large space for attachment of imaging targets. The construct with the shortest linker for each DARPin orientation was selected. In total, nine constructs were judged to be suitable for experimental characterization. These were based on the single DARPin noted earlier fused to one subunit of two different two-component cages, T33-21 (14) and T33-31 (32). Based on different helical lengths for connection to the DARPin, there were three candidate fusions to cage protein T33-31 and six fusions to cage protein T33-21.

### Cloning, Expression, and Purification

Constructs named DARP10, DARP11, DARP12, DARP14, and DARP16 were expressed and purified under conditions similar to those used for the cage proteins alone (14) with slight modifications. We purchased *E. coli* codon optimized gene fragments (Integrated DNA Technologies) and inserted the sequences encoding both cage subunit A and subunit B into a pET-22b vector, separated by the intergenic region of pET-DUET. Proteins were expressed in auto-induction media at 20 °C for two days.

Cells were suspended and lysed in Lysis Buffer (50mM Tris pH 8.0 250mM NaCl, 20mM imidazole) supplemented with DNase, lysozyme, and protease inhibitor (Thermo Scientific Pierce). Cleared lysate was loaded onto a HisTrap column (GE Healthcare) and eluted with a linear gradient of Elution Buffer (50mM Tris pH 8.0 250mM NaCl, 500mM imidazole). Pooled and concentrated fractions were then further purified with size-exclusion chromatography on a Superose 6 Increase column (GE Healthcare). Fractions corresponding to intact tetrahedral assemblies were used in further analysis.

### Negative stain electron microscopy

Freshly purified proteins at about 50 μg/mL were blotted onto glow-discharged 200 or 300 mesh copper formvar supported carbon grids (Ted Pella, Inc.), washed with Milli-Q water and stained with 2% uranyl acetate or 0.75% uranyl formate. Micrographs were collected using a Tecnai T12 with a bottom mount TVIPS F416 4K x 4K CMOS camera at a nominal magnification of 49,000x at the specimen level.

### Electron cryo-microscopy (cryo-EM)

#### DARP14 grid screening

Purified, concentrated DARP14 was screened for ice thickness, stability and particle distribution using a FEI TF20 microscope equipped with a bottom mount TVIPS F416, 4K x 4K CMOS camera.

#### DARP14 grid freezing for data collection

Superose 6 Increase column (GE Healthcare) purified, concentrated DARP14 was diluted to ~0.5 mg/mL using 10mM Tris pH 8.0, 500mM NaCl supplemented with 1mM of freshly prepared dithiothreitol (DTT) (Acros) and 4μL was pipetted on to C-Flat, carbon-coated, 1.2/1.3 200 Mesh copper grids (Electron Microscopy Sciences). Grids were blotted and frozen in liquid ethane using a Vitrobot Mark IV (FEI) and stored for data collection under liquid nitrogen.

#### Data Collection

Super-resolution movies were collected using a FEI titan krios (Thermo Fisher) microscope equipped with a Gatan K2 Summit direct electron detector at 22500X magnification at the specimen level with a physical pixel size of 1.31 Å/pixel (0.655 Å/pixel super-resolution).

### Data Processing

#### Cryo reconstructions

Super-resolution movies of frozen DARP14 were corrected for beam-induced motion using MotionCor2 (34). Particles were picked using the XMIPP software package. All co-ordinates were imported into and all micrographs analyzed using the RELION 2 software pipeline. An initial model used for all stages of 3D reconstructions was calculated *de-novo* using the Stochastic Gradient Descent algorithm in RELION 2.1-beta-0 using a subset of the calculated 2D classes. The final map containing DARPins was made while masking out all B-subunits. All calculations were done using different versions of RELION 2 except for the final refinements, post-processing and local resolution estimations (including the final fourier shell correlation (FSC) calculations), which were done using RELION version 2.1-beta-1. Masks were created from the reconstructions using combinations of both RELION and UCSF Chimera.

#### Structure Analysis

All reconstructions were analyzed using UCSF Chimera. The design model was initially fit using UCSF Chimera followed by refinements using Phenix and Rosetta. Refined models were analyzed using UCSF Chimera, PyMOL and COOT.

## Acknowledgements

The authors thank Marianne Vo for her contribution to protein production, James Evans for preliminary assistance with the EM studies and Johan Hattne (Gonen Lab HHMI Janelia) for help with the computational cluster. This work was funded by the BER program of the Department of Energy Office of Science, award DE-FC02-2ER63421, and by the UCLA Whitcome Fellowship to YL. The Gonen laboratory is supported by the Howard Hughes Medical Institute.

## Author Contributions

TOY helped design and supervise the research. YL performed the protein design and characterization experiments. SG and TG performed and analyzed the cryo-electron microscopy studies. All authors contributed to writing the manuscript.

## Data Deposition

The averaged EM density map will be deposited in the EMDB. The molecular model of the cage plus DARPin assembly fit into the EM map will be deposited in the PDB.

## Supplementary Text

### Protein Sequences

Protein sequences of soluble genetic fusions between protein cages and the chosen DARPin. The regions corresponding to the DARPins are underlined. Additional residues designed between the cage subunit and its joining DARPin are double underlined.

- DARP10: Subunit A: MEEVVLITVPSALVAVKIAHALVEERLAACVNIVPGLTSIYREEGSVVSDH ELLLLVKTTTDAFPKLKERVKELHPYEVPEIVALPIAEGNREYLDWLREN <u>MERARQELGKKLLEAARAGQDDEVRILMANGADVNAHDDQGSTPLHLAAWIGHPEIVEVLLKHGADVNARDTDGWTPLHLAADNGHLEIVEVLLKYGADVNAQDAYGLTPLHLAADRGHLEIVEVLLKHGADVNAQDKFGKTAFDISIDNGNEDLAEILQKLN</u> Subunit B: MVRGIRGAITVEEDTPAAILAATIELLLKMLEANGIQSYEELAAVIFTVTED LTSAFPAEAARLIGMHRVPLLSAREVPVPGSLPRVIRVLALWNTDTPQDR VRHVYLNEAVRLRPDLESAQLEHHHHHH
- DARP11 Subunit A: MRITTKVGDKGSTRLFGGEEVWKDSPIIEANGTLDELTSFIGEAKHYVDEE MKGILEEIQNDIYKIMGEIGSKGKIEGISEERIAWLLKLILRYMEMVNLKSF VLPGGTLESAKLDVCRTIARRALRKVLTVTREFGIGAEAAAYLLALSDLLF LLARVIEIEKN<u>DLGKKLLEAARAGQDDEVRILMANGADVNAHDDQGSTPLHLAAWIGHPEIVEVLLKHGADVNARDTDGWTPLHLAADNGHLEIVEVLLKYGADVNAQDAYGLTPLHLAADRGHLEIVEVLLKHGADVNAQDKFGKTAFDISIDNGNEDLAEILQKLN</u> Subunit B: MPHLVIEATANLRLETSPGELLEQANKALFASGQFGEADIKSRFVTLEAYR QGTAAVERAYLHACLSILDGRDIATRTLLGASLCAVLAEAVAGGGEEGV QVSVEVREMERLSYAKRVVARQRLEHHHHHH
- DARP12 Subunit A: MRITTKVGDKGSTRLFGGEEVWKDSPIIEANGTLDELTSFIGEAKHYVDEE MKGILEEIQNDIYKIMGEIGSKGKIEGISEERIAWLLKLILRYMEMVNLKSF VLPGGTLESAKLDVCRTIARRALRKVLTVTREFGIGAEAAAYLLALSDLLF LLARVIEIEKN<u>QDLGKKLLEAARAGQDDEVRILMANGADVNAHDDQGSTPLHLAAWIGHPEIVEVLLKHGADVNARDTDGWTPLHLAADNGHLEIVEVLLKYGADVNAQDAYGLTPLHLAADRGHLEIVEVLLKHGADVNAQDKFGKTAFDISIDNGNEDLAEILQKLN</u> Subunit B: MPHLVIEATANLRLETSPGELLEQANKALFASGQFGEADIKSRFVTLEAYR QGTAAVERAYLHACLSILDGRDIATRTLLGASLCAVLAEAVAGGGEEGV QVSVEVREMERLSYAKRVVARQRLEHHHHHH
- DARP14: Subunit A: MRITTKVGDKGSTRLFGGEEVWKDSPIIEANGTLDELTSFIGEAKHYVDEE MKGILEEIQNDIYKIMGEIGSKGKIEGISEERIAWLLKLILRYMEMVNLKSF VLPGGTLESAKLDVCRTIARRALRKVLTVTREFGIGAEAAAYLLALSDLLF LLARVIEIE<u>LGKKLLEAARAGQDDEVRILMANGADVNAHDDQGSTPLHLAAWIGHPEIVEVLLKHGADVNARDTDGWTPLHLAADNGHLEIVEVLLKYGADVNAQDAYGLTPLHLAADRGHLEIVEVLLKHGADVNAQDKFGKTAFDISIDNGNEDLAEILQKLN</u> Subunit B: MPHLVIEATANLRLETSPGELLEQANKALFASGQFGEADIKSRFVTLEAYRQGTAAVERAYLHACLSILDGRDIATRTLLGASLCAVLAEAVAGGGEEGVQVSVEVREMERLSYAKRVVARQRLEHHHHHH
- DARP16 Subunit A: MRITTKVGDKGSTRLFGGEEVWKDSPIIEANGTLDELTSFIGEAKHYVDEE MKGILEEIQNDIYKIMGEIGSKGKIEGISEERIAWLLKLILRYMEMVNLKSF VLPGGTLESAKLDVCRTIARRALRKVLTVTREFGIGAEAAAYLLALSDLLF LLARVIEIE<u>DLGKKLLEAARAGQDDEVRILMANGADVNAHDDQGSTPLHLAAWIGHPEIVEVLLKHGADVNARDTDGWTPLHLAADNGHLEIVEVLLKYGADVNAQDAYGLTPLHLAADRGHLEIVEVLLKHGADVNAQDKFGKTAFDISIDNGNEDLAEILQKLN</u> Subunit B: MPHLVIEATANLRLETSPGELLEQANKALFASGQFGEADIKSRFVTLEAYRQGTAAVERAYLHACLSILDGRDIATRTLLGASLCAVLAEAVAGGGEEGVQVSVEVREMERLSYAKRVVARQRLEHHHHHH

**SI Fig. 1.**
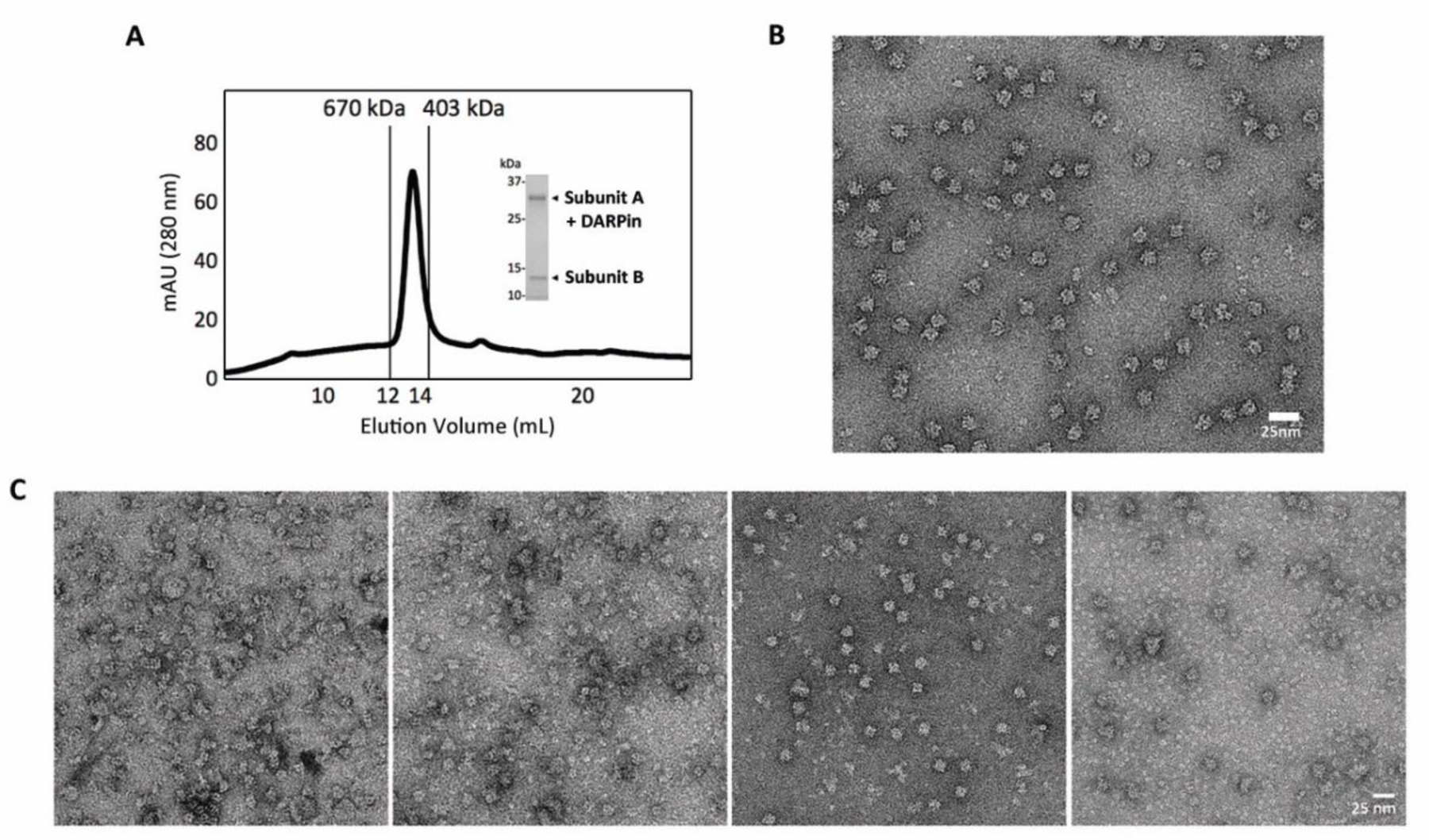
Designed DARPin-displaying cages form particles of expected size and shape. A. Purification of the DARPin14 cage scaffold indicates a homogeneous preparation by size exclusion chromatography, SDS PAGE analysis and negative stain electron microscopy. The two components in DARP14 co-elutes and migrate as single peak with the correct retention volume from size-exclusion chromatography (Superose 6 Increase, GE Healthcare). B. Negative stain EM of DARP14. C. Negative stain EM of DARP10, DARP11, DARP12, and DARP16, from left to right. Particles of ~15nm diameter are clearly visible.

**SI Fig. 2.**
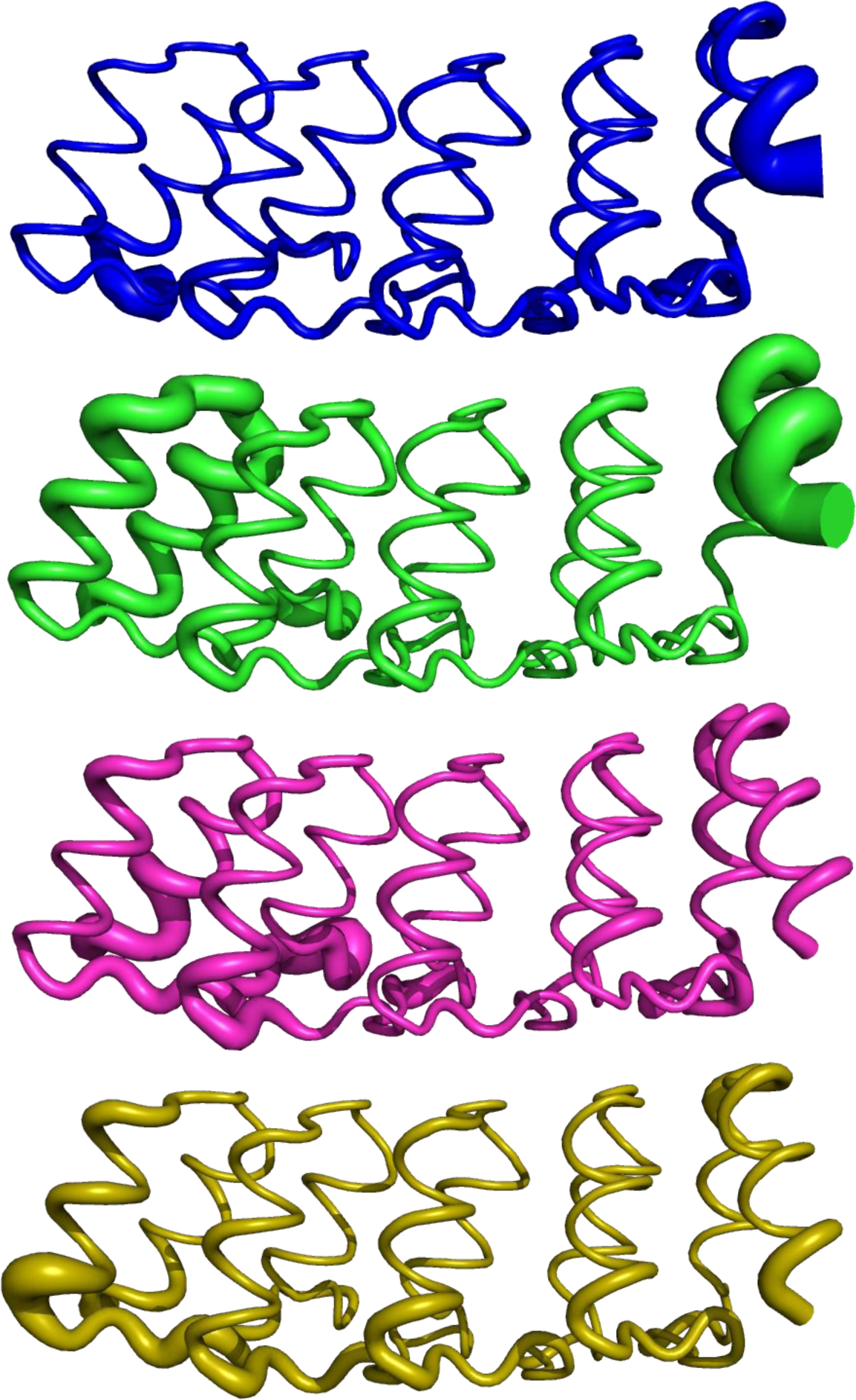
Comparison of thermal atomic displacement parameters (B-factors) from previous DARPin crystal structures. Structures of different DARPins shown in sausage representation according to their B-factors. Structures are positioned in similar orientation with N-termini on the left and C-termini on the right. Top to bottom: consensus DARPin (1MJ0), DARPin specific for ERK2 binding as used here in DARP14 (ERK2 omitted for clarity, 3ZU7), consensus DARPins with stabilizing mutations on the C-terminal repeat (2XEE & 2XEH). The C-terminal repeat tends to have higher B-factor than other repeats, unless stabilized by additional mutations.

**SI Table 1.**
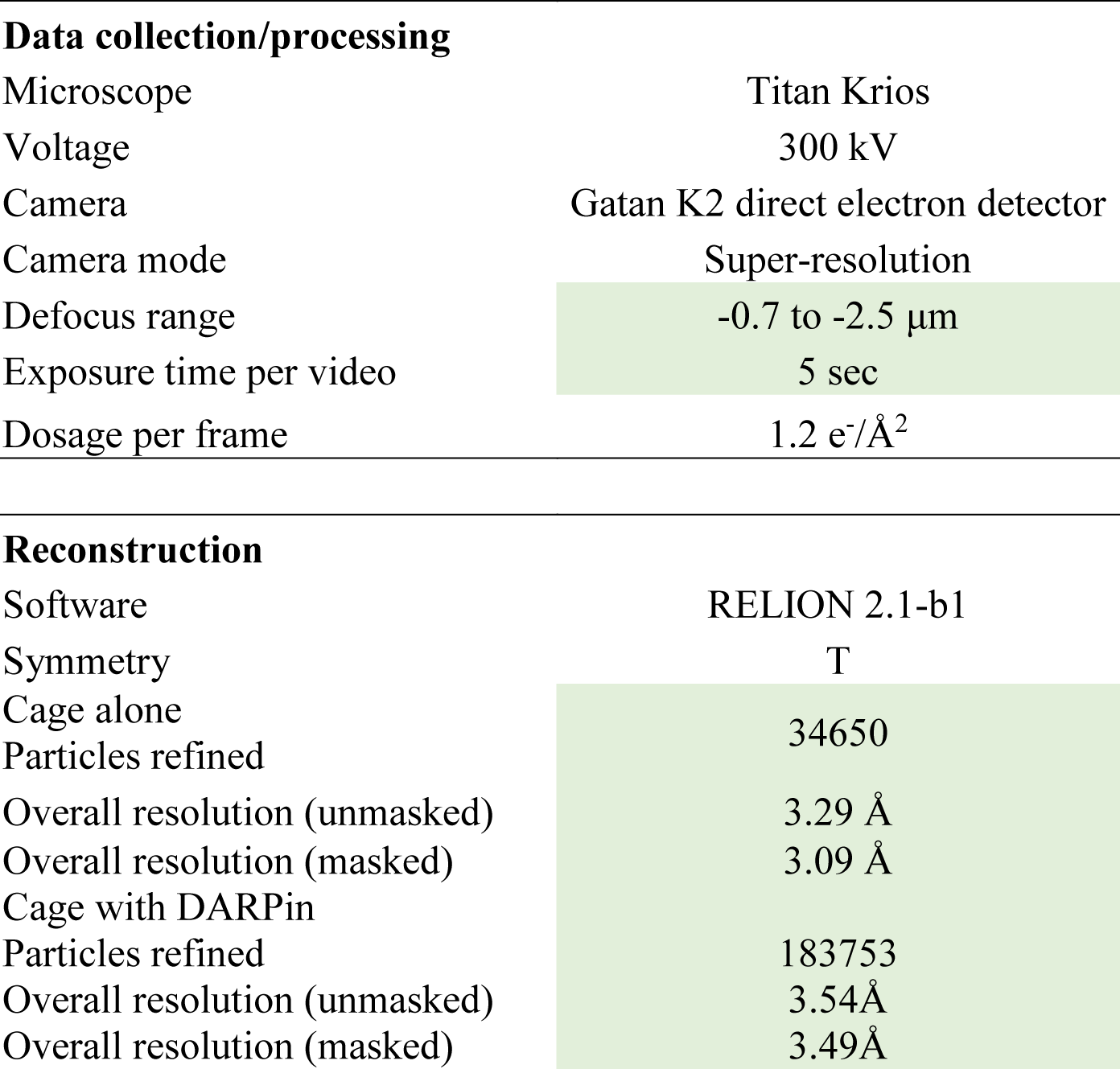
CryoEM data table

